# Self-supervised learning on millions of pre-mRNA sequences improves sequence-based RNA splicing prediction

**DOI:** 10.1101/2023.01.31.526427

**Authors:** Ken Chen, Yue Zhou, Maolin Ding, Yu Wang, Zhixiang Ren, Yuedong Yang

**Affiliations:** School of Computer Science and Engineering, Sun Yat-sen University, Guangzhou, China; Peng Cheng Laboratory, Shenzhen, China; Key Laboratory of Machine Intelligence and Advanced Computing (Sun Yat-sen University), Ministry of Education, China

## Abstract

RNA splicing is an important post-transcriptional process of gene expression in eukaryotic cells. Predicting RNA splicing from primary sequences can facilitate the interpretation of genomic variants. In this study, we developed a novel self-supervised pre-trained language model, SpliceBERT, to improve sequence-based RNA splicing prediction. Pre-training on pre-mRNA sequences from vertebrates enables SpliceBERT to capture evolutionary conservation information and characterize the unique property of splice sites. SpliceBERT also improves zero-shot prediction of variant effects on splicing by considering sequence context information, and achieves superior performance for predicting branchpoint in the human genome and splice sites across species. Our study highlighted the importance of pre-training genomic language models on a diverse range of species and suggested that pre-trained language models were promising for deciphering the sequence logic of RNA splicing.

## INTRODUCTION

Ribonucleic acids (RNA) splicing is a fundamental post-transcriptional process in eukaryotic gene expression, which removes introns from precursor messenger RNAs (pre-mRNAs) and ligates exons into mature mRNAs. Though the mechanism underlying RNA splicing is complex, a variety of studies have found that some key determinants of splicing are encoded in DNA sequences^1–3^. Therefore, deciphering splicing codes from RNA sequences by computational models is a promising approach and will facilitate the interpretation of genetic variants that affect RNA splicing^4^.

Early studies mainly aimed to identify short sequence motifs related to splicing, including exonic splicing enhancers^5^, branchpoints^6^ and other splicing factors^7^ with statistical models. Benefiting from accumulated high-throughput sequencing data, subsequent studies were able to employ machine learning and deep learning models to directly predict RNA splicing events from primary sequences. For example, HAL^8^ is a model trained on alternative splicing events from millions of random sequences for predicting the change of exon skipping and 5’ alternative splicing induced by genetic variants. SPANR^3^ is a Bayesian network model for predicting the Percent Spliced In (PSI or Ψ) of alternatively spliced exons. More recently, deep learning models like MMSplice^9^, SpliceAI^10^ and Pangolin^11^ employed deep convolutional neural networks (CNN) to predict alternative splicing events, splice sites or splice site usage. These methods achieved superior performance as compared to earlier studies and have been widely utilized to analyze aberrant RNA splicing events caused by genetic variants. Though significant progress has been made in this field, there is still room for further exploration. For instance, state-of-the-art splicing models were developed to predict splice sites or alternative splicing from primary sequences, while evolutionary information was not considered. Moreover, another important splicing regulator, branchpoint, was less studied. Unlike splice sites, which can be reliably detected from RNA-seq data ^12,13^, BPs are much more difficult to be identified^14^, making it challenging to develop deep learning models for BP prediction due to the lack of adequate high-confidence BPs.

To alleviate the problem of insufficient data, the self-supervised learning (SSL) method utilized by large pre-trained language models (pLMs)^15–17^ can be adopted. A common form of SSL is masked language modeling (MLM) and it has already been adopted to develop pLMs of protein^18,19^, non-coding RNA (ncRNA)^20^ and prokaryote genome^21^ sequences. These models were pre-trained on a large number of sequences from a diverse range of species, and thereby captured the evolutionary information which is critical for sequence-based modeling. However, these models cannot be directly applied to RNA splicing because eukaryotic protein-coding RNA sequences are very different from ncRNAs or prokaryote genome sequences. Though there are genome pLMs^22,23^ like DNABERT^24^, they were pre-trained on only the human genome, and thus it remains unclear whether pLMs pre-trained on sequences from more species could improve sequence-based RNA splicing prediction.

Here, we developed a novel pre-trained model, SpliceBERT, for studying RNA splicing. SpliceBERT was pre-trained by masked language modeling on over 2 million precursor messenger RNA (pre-mRNA) sequences from 72 vertebrates. Compared to pLMs for only the human genome, SpliceBERT can more effectively capture evolutionary conservation from primary sequences. The hidden states and attention weights generated by SpliceBERT can reflect the functional characteristics of splice sites. SpliceBERT also improved zero-shot prediction of variant effects on RNA splicing by incorporating context information. These findings suggest that self-supervised learning is beneficial to learn biologically meaningful representations from sequences. On branchpoint and cross-species splice site prediction, SpliceBERT exhibited superior performance to conventional baseline models and other human-only pLMs, further indicating the effectiveness of pre-training models on a diverse range of species. The SpliceBERT model is available at https://github.com/biomed-AI/SpliceBERT.

## RESULTS

### Masked language modeling captures evolutionary conservation information

SpliceBERT is a pre-trained pre-mRNA language model (LM) for studying RNA splicing (Figure 1A). It was developed based on the Bidirectional Encoder Representations from Transformers (BERT)^16,25^ architecture and pre-trained by masked language modeling (MLM) on over 2 million pre-mRNA sequences from 72 vertebrates (see Methods). For MLM pre-training, SpliceBERT was set to predict the type of nucleotides that were masked in the input sequences, by which it could learn dependencies between nucleotides in a self-supervised manner. Figure 1B illustrates that the balanced accuracy (ACC) for nucleotide type prediction in MLM is 0.641 (0.636) and 0.493 (0.511) for introns and exons, respectively, in coding (non-coding) genes. The higher ACC in introns is likely attributed to its higher sequence repeat contents: 46.1%/51.3% in coding/non-coding genes versus 2.4%/37.1% in coding/non-coding genes (Figure 1C). Similar results can be found in different functional regions when repetitive and non-repetitive sequences were analyzed separately, indicating that the logic of non-coding sequences is easier to be captured than coding sequences due to the enrichment of repetitive sequences.

**Figure 1.**
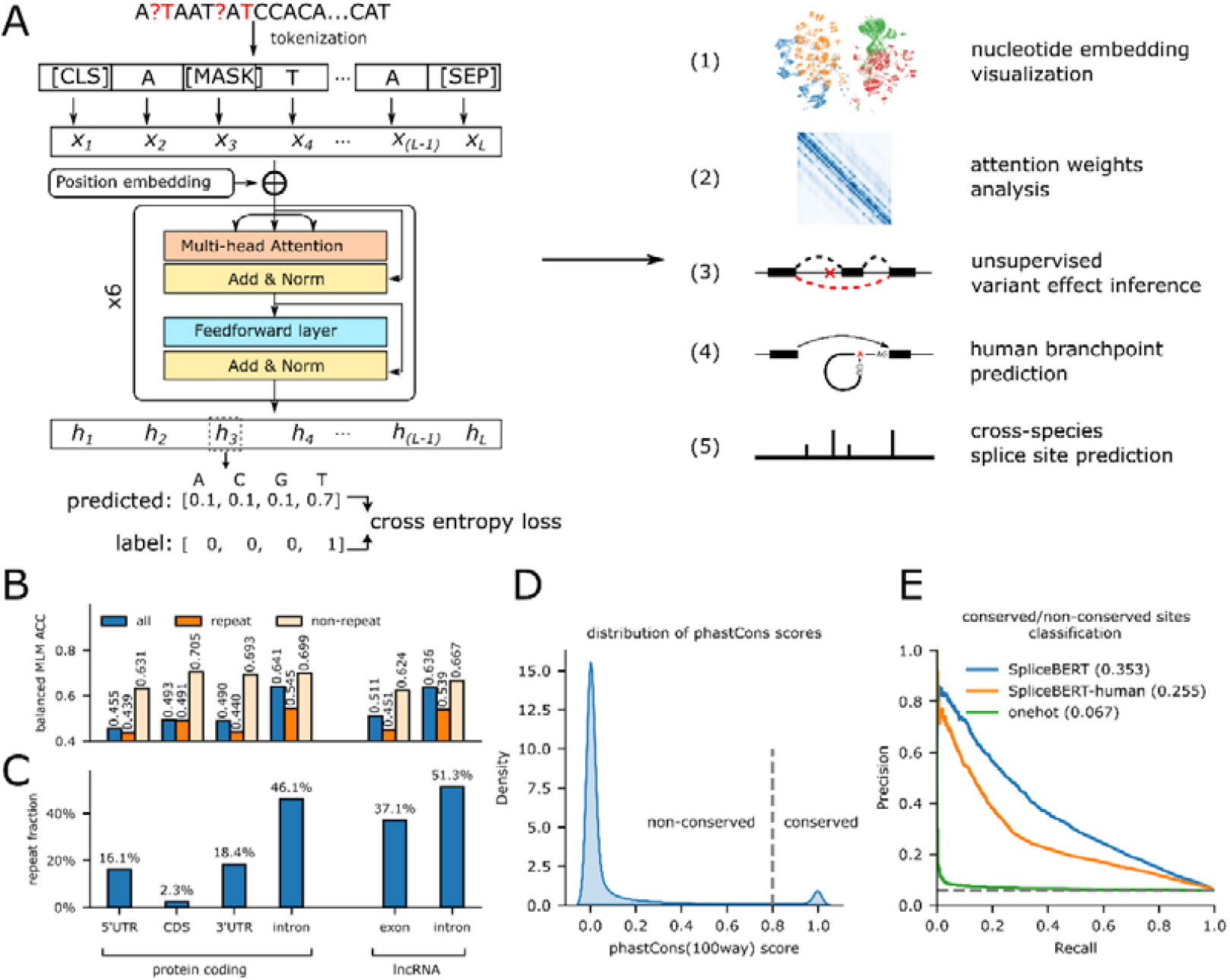
Pre-training SpliceBERT on pre-mRNA sequences by masked language modeling. (**A)** SpliceBERT is consisted of 6 Transformer encoder layers and was pre-trained on pre-mRNA sequences of vertebrates. It is applied to analyze the learned nucleotide representations, identify splice-disrupting variants, predict branchpoint sites and predict splice sites. (**B**) Balanced accuracy of masked token prediction (MLM ACC) in repetitive/non-repetitive regions of different functional genomic regions. (**C**) The fraction of repeats in different functional regions of protein-coding and lncRNA transcripts. (**D**) The distribution of phastCons(100way) score in transcripts. The cut-off of conserved/non-conserved sites is set to 0.8. (**E**) The precision-recall curves of logistic regression models for distinguishing between nucleotides at conserved and non-conserved sites based on nucleotide embeddings from SpliceBERT, SpliceBERT-human and one-hot encoding. (UTR: untranslated regions, CDS: coding sequences, lncRNA: long non-coding RNA)

It is of interest to investigate whether MLM pre-training could enable SpliceBERT to capture evolutionary information, because most methods for identifying evolutionarily conserved elements are based on multiple sequence alignment (MSA) ^26^, which is time-consuming. To this end, we extracted hidden states in the last encoder layer of SpliceBERT and fed them to a logistic regression (LR) model to see if they can be exploited to distinguish between conserved (phastCons ≥ 0.8) and non-conserved (phastCons < 0.8) sites (Figure 1D) (see Methods). As shown in Figure 1E, the SpliceBERT-based LR model achieved an average precision (AP) score of 0.353, outperforming the baseline models that rely on SpliceBERT-human (AP=0.255) and one-hot encoded (AP=0.067) embeddings. This demonstrated that MLM pre-training is able to capture evolutionary information, and this ability can be enhanced by augmenting the pre-training data with sequences from more species. To be noted, the AP score was merely used to compare the performance of different models, rather than measuring their absolute ability. This is because the phastCons scores were derived from 100 species while our model was pre-trained on only 72 species. The ability of SpliceBERT for capturing evolutionary information is potentially beneficial for sequence-based modeling tasks and we thus conducted further analysis in the following sections to illustrate this point.

### Nucleotide embeddings learned by SpliceBERT characterize the property of splice sites

The nucleotide embeddings learned by SpliceBERT were studied to assess whether they were able to characterize the biological properties of splice sites. To achieve this, canonical splice sites (SS) and non-splice GT/AG sites (NSS) were collected from the human genome and the nucleotide embeddings of them were generated using SpliceBERT and 3 baseline methods (SpliceBERT-human, DNABERT and one-hot). These embeddings were visualized by the UMAP^27^ algorithm and clustered by the Leiden^28^ algorithm (see Methods). As shown in Figure 2A, SpliceBERT achieved the highest normalized mutation information (NMI) score (NMI=0.31/0.31 for GT/AG, respectively) for distinguishing between SS and NS, surpassing SpliceBERT-human (NMI=0.18/0.08), DNABERT (NMI=0.05/0.02) and one-hot encoding (NMI=0.08/0.06). Though DNABERT and SpliceBERT-human were both pre-trained on the human data, the lower NMI of DNABERT may stem from its k-mer tokenization strategy, which tends to generate more clusters as the value of k increases (Supplementary Figure S1). Next, we conducted the same analysis on splice sites of high (SSE > 0.8) and low (SSE < 0.2) strength estimated by SpliSER^29^ in the K562 cell line. This presents a more challenging scenario as all the samples are authentic splice sites. As expected, the NMI scores decreased for all the models (Figure 2B), while SpliceBERT still achieved the highest NMI score (NMI=0.20/0.21 for donor/acceptor). These findings indicated that SpliceBERT trained on a variety of species is more powerful to capture the sequence determinants of splice sites than human-only pLMs.

**Figure 2.**
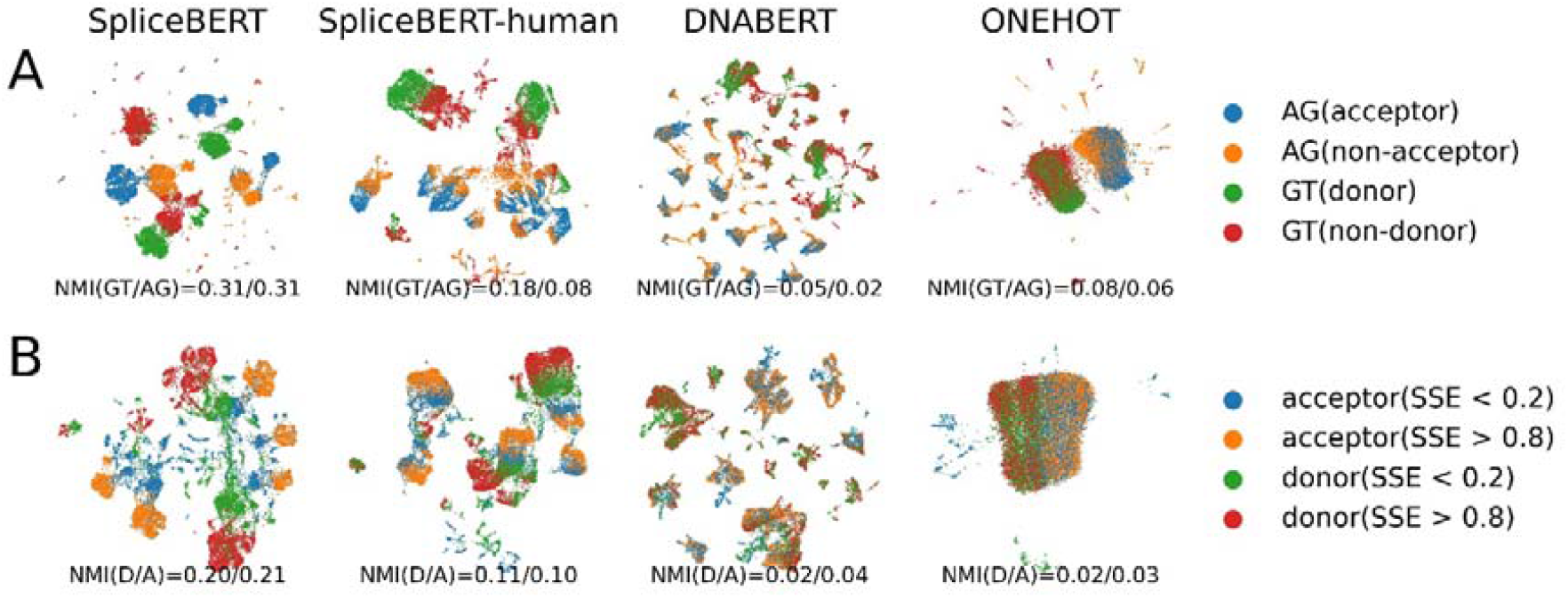
Investigating the nucleotide embeddings of splice sites. UMAP visualization of nucleotide embeddings generated by different methods for (**A**) canonical splice sites (GT/AG) and non-splice GT/AG sites and (**B**) splice sites of high and low usage in K562 estimated by SpliSER. (NMI: normalized mutual information. D: donor. A: acceptor)

In addition to the last layer in SpliceBERT, we found that layers 2-5 are also informative for distinguishing SS from NSS (Supplementary Figure S3). The optimal performance was achieved by hidden states in the 4^th^ layer but not the last layer (the 6^th^ layer). This is probably because the hidden states of the last layer are mostly related to predicting the masked tokens during pre-training^30^, while the intermediate layers preserve more contextual information.

### Attention weights in SpliceBERT correlate with donor-acceptor dependencies

We next investigated the attention weights in SpliceBERT for the functional associations between splice donor and acceptor sites. Concretely, the attention weights between different donor/acceptor combinations (Supplementary Figure S4) or randomly selected nucleotides were extracted from the multi-head attention module in SpliceBERT (see Methods). The attention weights between donor or acceptor pairs were consistently higher than those between random site pairs (Figure 3A), which was likely due to the higher evolutionary conservation around splice sites than the other sites (Figure 3B). More importantly, donor-acceptor pairs from the same introns achieved the highest attention weights among all the groups, including those from the same exons (P-value < 10^−6^, by Mann-Whitney test). This implied that SpliceBERT captured the close functional association between donors and acceptors from the same introns, which is in line with the intron-centric nature of RNA splicing ^31,32^ as well as the enrichment of conserved complementary regions in the ends of introns^33^. To investigate the contribution of different encoder layers, we analyzed the attention weights between donor-acceptor pairs from the same introns from layer 1 to layer 6, respectively. The attention weights of the donor (acceptor) sites were aligned with respect to the acceptor (donor) sites and the median values across all the samples were calculated. As illustrated in Figure 3C, peaks of attention weights can be observed around the acceptor/donor sites in the 3^rd^, 4^th^, and 5^th^ layer, which is 1.9, 16.1 and 1.7 times that of attention weights averaged across the entire regions (Supplementary Table S1). In contrast, the weights of donors at acceptors are only 0.7, 0.5 and 0.7 times the average attention in the 1^st^, 2^nd^ and 6^th^ layer, respectively. This indicates that the attention weights in layers 3-5 can better capture donor-acceptor associations, which is consistent with our observation in the analysis of hidden states (Supplementary Figure S3). Taken together, we could conclude that the attention weights in SpliceBERT can reflect the functional associations between splice donors and acceptors.

**Figure 3.**
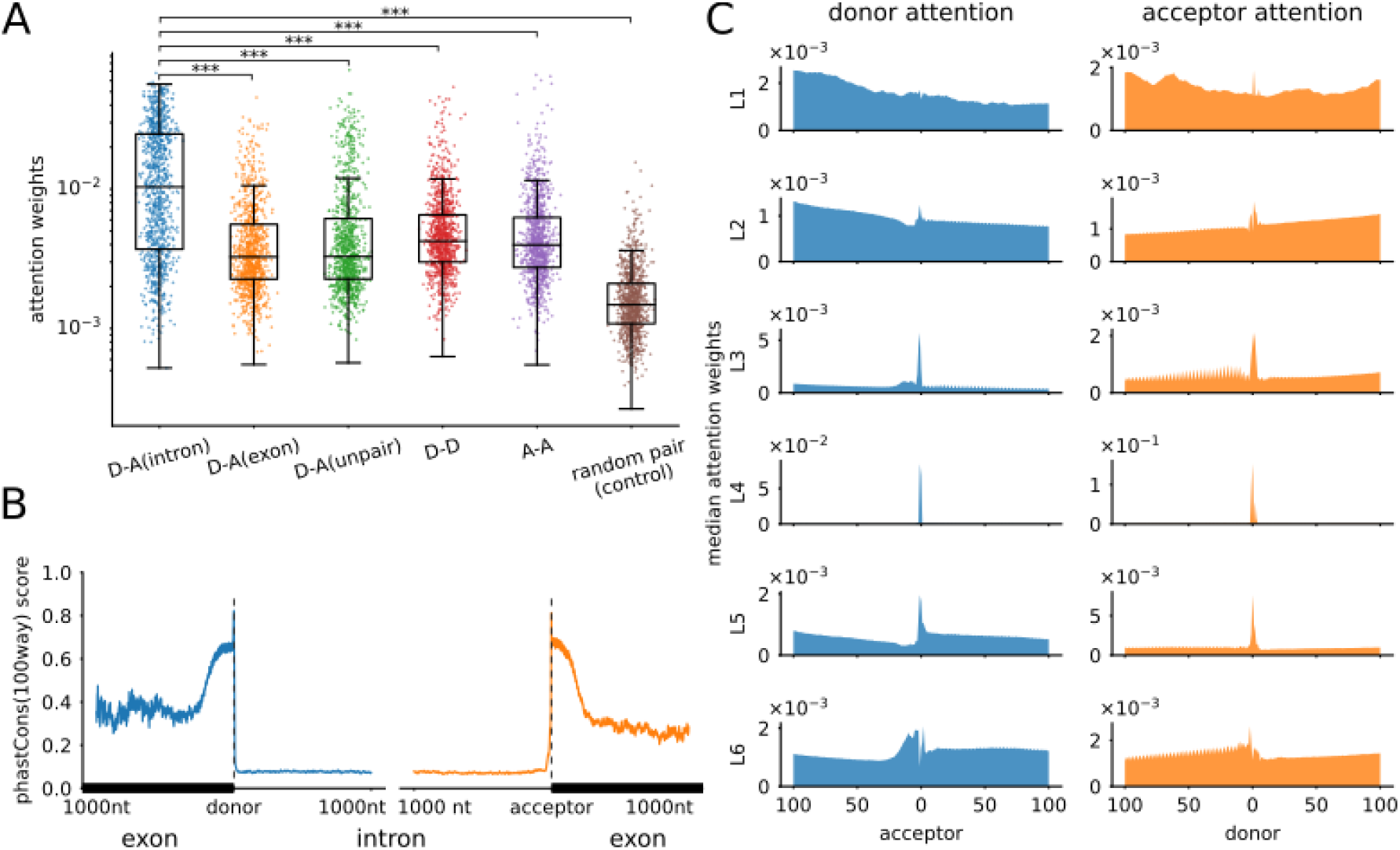
Analyzing the attention weights in SpliceBERT. **(A)** The distribution of attention weights between donors and acceptors within the same intron (D-A(intron)), within the same exon (D-A(exon)), across different intron/exon (D-A(unpair)), among donors (D-D) and among acceptors (A-A). A control group of randomly paired nucleotides is also shown (random pair). The average attention weights across all the 6 layers in SpliceBERT were used. The statistical significance was assessed by *t*-test. **(B)** The distribution of phastCons(100way) scores around donors and acceptors within 1000nt (from the same introns). **(C)** The distribution of donors’ and acceptor’s attention weights around acceptors and donors, respectively, in each Transformer layer. The median attention weights across different samples are shown. (D-A: donor-acceptor, D-D: donor-donor, A-A: acceptor-acceptor, L: layer. ^***^: P-value <)

### SpliceBERT-derived context information improves zero-shot variant effects prediction

To apply SpliceBERT for predicting the effects of variants on RNA splicing in a zero-shot fashion, we measured the influence of single nucleotide variants (SNVs) on adjacent nucleotides by calculating the KL-divergence between normalized MLM logits of wild-type (WT) and mutant (MT) sequences (see Methods). On two datasets of variants related to splicing, MFASS^34^ and Vex-seq^35^, splice-disrupting variants (SDVs) tended to have a larger impact on the MLM logits of adjacent nucleotides than non-SDVs (Figure 4A and 4B). More generally, we also studied the variants at conserved (phastCons ≥ 0.8) and non-conserved (phastCons < 0.8) sites because evolutionarily conserved regions in genomic sequences were usually with critical functions. As expected, the variants at conserved sites tend to induce greater change in MLM logits than those at non-conserved sites (Figure 4C), further indicating that MLM logits of non-mutated sites can be exploited to indicate the functional effects of genetic variants.

**Figure 4.**
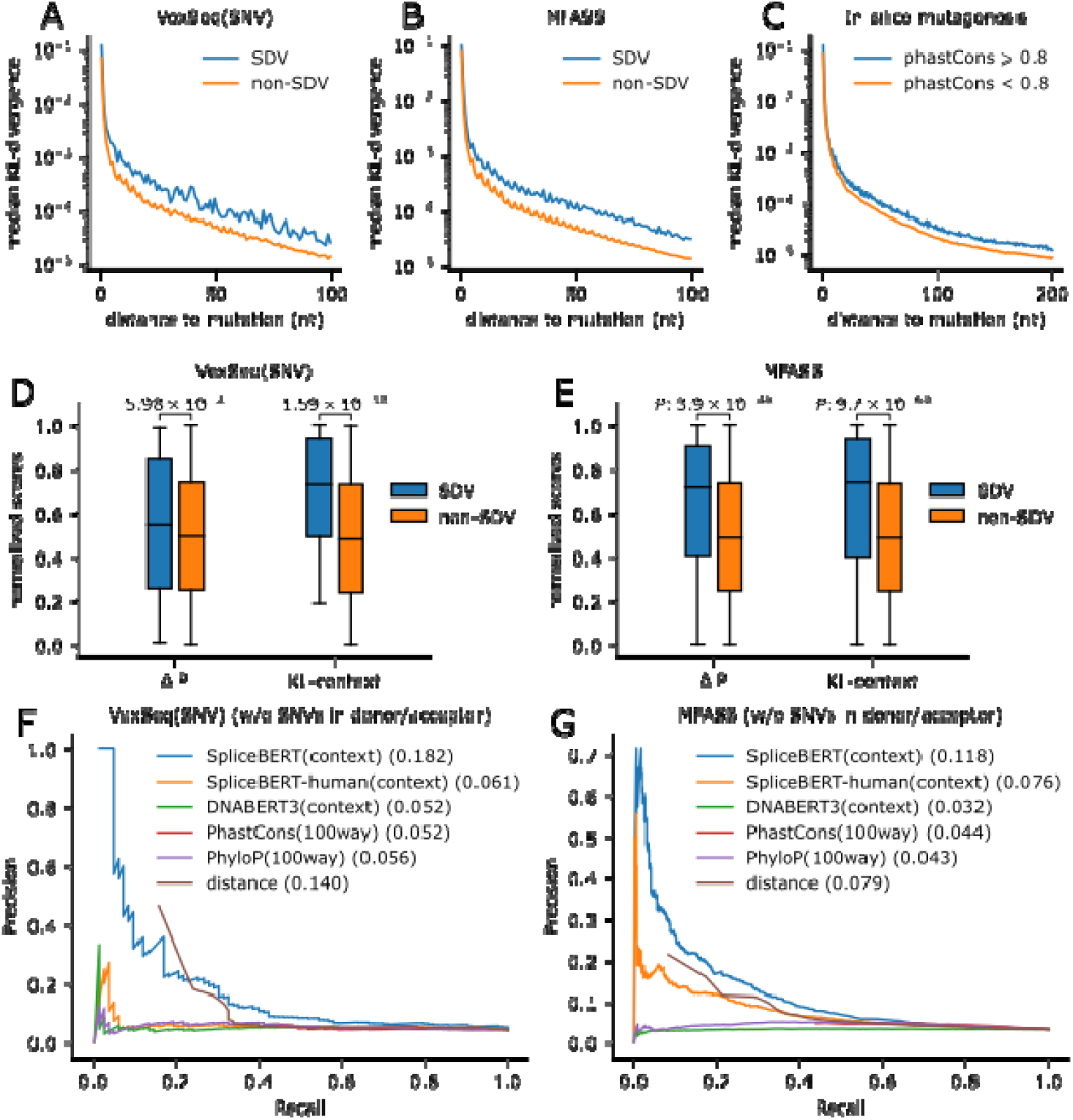
Applying SpliceBERT to zero-shot prediction of variant effects on RNA splicing. The median value of KL-divergence between the MLM logits of wild-type and mutant sequences for splice-disrupting (SDV) and non-splice-disrupting (non-SDV) variants in **(A)** Vex-seq and (**B**) MFASS. **(C)** The median value of KL-divergence around variants at conserved and non-conserved sites. The boxplot of delta-logit (P) and KL-context score of SDV and non-SDV group in (**D**) Vex-seq and (**E**) MFASS. The precision-recall curves of SpliceBERT and baseline methods for classifying SDV and non-SDV samples in (**F**) Vex-seq and (**G**) MFASS.

Therefore, we took the sum of KL-divergence (in logarithm scale) within 100nt up- and downstream of each variant (KL-context score) for inferring the effects on RNA splicing. Compared with the Δ*P*, metric adopted by previous studies^36,37^, which measures the change of allele logit, the difference in median KL-context score of SDVs and non-SDVs is larger in Vex-seq and MFASS (Figure 4D and 4E), respectively. Next, we quantified the performance of KL-context scores derived from different pLMs (SpliceBERT, SpliceBERT-human and DNABERT), phastCons, phyloP and distance to splice sites on Vex-seq and MFASS, while excluding the SNVs at donor/acceptor sites due to their high prevalence as SDVs (Vex-seq: 68.2%, MFASS: 80.7%, Supplementary Figure S5). As shown in Figure 4F and 4G, SpliceBERT-derived KL-context scores consistently outperformed human-only pLMs (SpliceBERT-human and DNABERT), conservation scores (phyloP and phastCons) and the distance from variants to splice sites. These results demonstrated that incorporating the context information derived from SpliceBERT can improve zero-shot inference of variant effects on RNA splicing.

### SpliceBERT improves branchpoint prediction

In addition to the splice site, branchpoint (BP) is another crucial type of splicing regulator, which involves splice acceptor identification^38^. We first generated the embeddings of BPs and non-BP sites that conform to the typical YTNAY motif^14^ and visualized them by UMAP (see Methods). The BP embeddings generated by SpliceBERT achieved an NMI score of 0.090 by Leiden clustering (Figure 5A), which is much lower than that of the splice sites (Figure 2). The performance of DNABERT and one-hot encoding is even worse (NMI = 0.084/0.087). This indicates that BPs are difficult to be characterized solely through MLM pre-training, which is possible because BP sequences are highly degenerate^14^ and are usually challenging to be accurately predicted based on sequences^38^.

**Figure 5.**
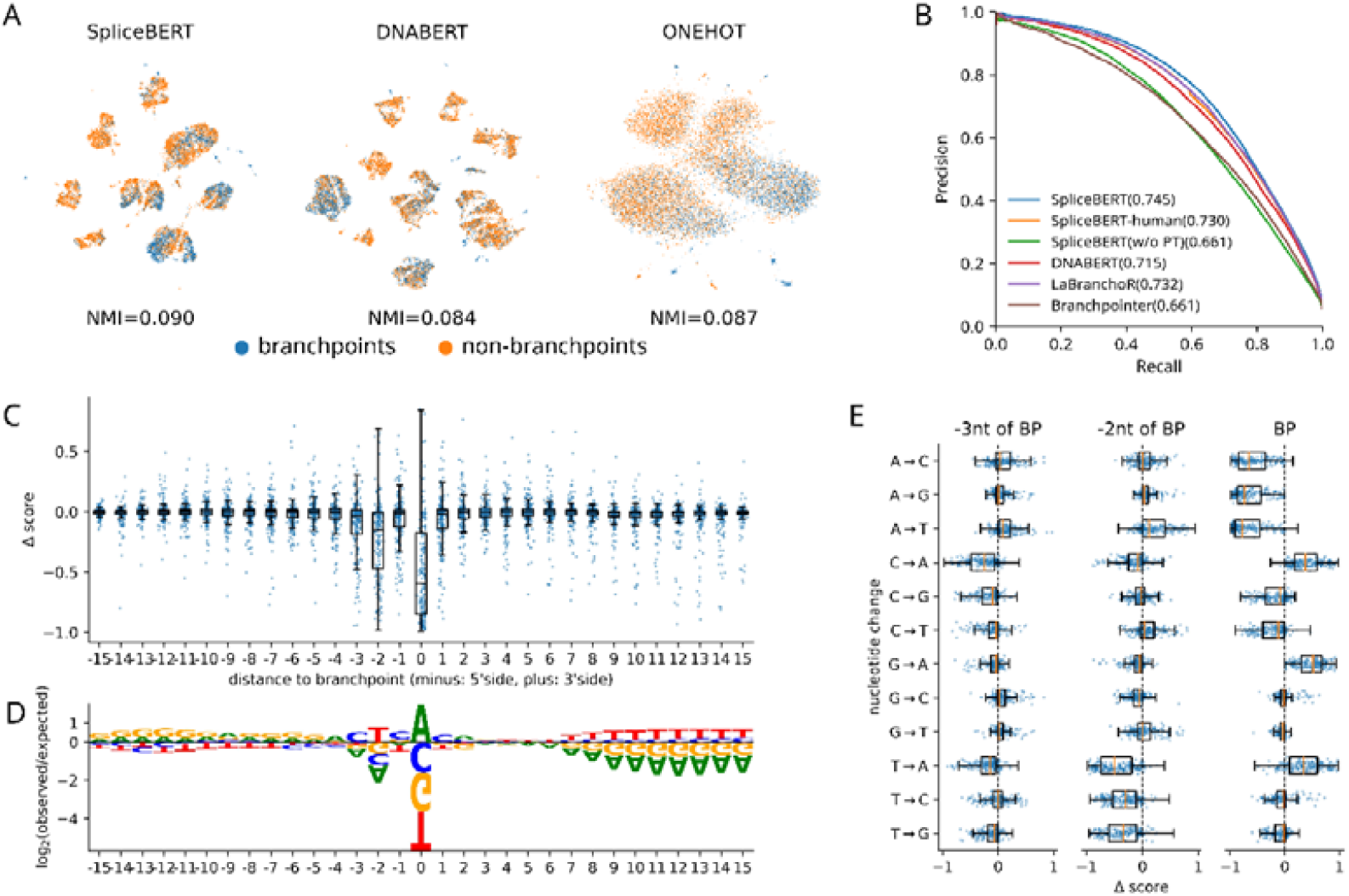
Predicting human branchpoint sites using SpliceBERT. (**A**) The UMAP visualization of branchpoint and non-branchpoint site embeddings. The normalized mutual information (NMI) scores were calculated and averaged for each sequence motif, respectively. (**B**) The precision-recall curves of the models for predicting branchpoints. (**C**) The impact of variants at different sites on SpliceBERT predicted branchpoint scores and (**D**) the sequence consensus around branchpoints. (**E**) The predicted change of branchpoint (BP) score induced by different mutation types at the BP sites and 2nt- and 3nt-upstream of BP sites.

To enhance sequence-based BP prediction, SpliceBERT was finetuned on the Mercer’s dataset that includes branchpoints identified in the human genome (see Methods). We adopted the 10-fold nested cross-validation strategy to avoid over-fitting and make full use of the samples for evaluation (see Methods). SpliceBERT achieved an AP score of 0.745, outperforming SpliceBERT-human, DNABERT, LaBranchoR^39^, and Branchpointer^40^ by 2.1%, 4.2%, 1.8% and 12.7%, respectively (Figure 5B). Additionally, the pre-training process is found to be indispensable, as the AP score of the SpliceBERT model trained from scratch without pre-training (SpliceBERT w/o PT) is only 0.661. These results demonstrated that SpliceBERT can improve sequence-based prediction of human branchpoints. In silico mutagenesis indicated that the variants that have a significant impact on branchpoint prediction enriched within the range of 3nt upstream to 1nt downstream of the BP site ([BP-3, BP+1]), consistent with the known BP motif (Figure 5C and 5D). The loss of adenine at BP sites significantly reduced predicted BP scores, and variant T>A and T>G at BP-2 can also largely decrease predicted BP scores (Figure 5E). These observations are consistent with the known patterns of pathogenic variants around BP sites as reported in Zhang et al’s study^41^.

### SpliceBERT improves cross-species splice site prediction

SpliceBERT was expected to achieve better performance in cross-species prediction since it was pre-trained on more than 70 vertebrates. To address this, we finetuned SpliceBERT to predict splice site on the Spliceator^42^ training dataset, which included splice sites in over 100 species, and tested it on test datasets from zebrafish, fruit fly, worm and arabidopsis (see Methods). As illustrated in Figure 6A, SpliceBERT achieved superior performance to baseline models on the test datasets (P-value < 0.05, by paired *t*-test, Supplementary Table S2). In particular, SpliceBERT surpassed SpliceBERT-human and DNABERT by an average of 1.3% and 2.1%, respectively. This suggests that SpliceBERT compares favorably to the models pre-trained on only human sequences in cross-species prediction.

**Figure 6.**
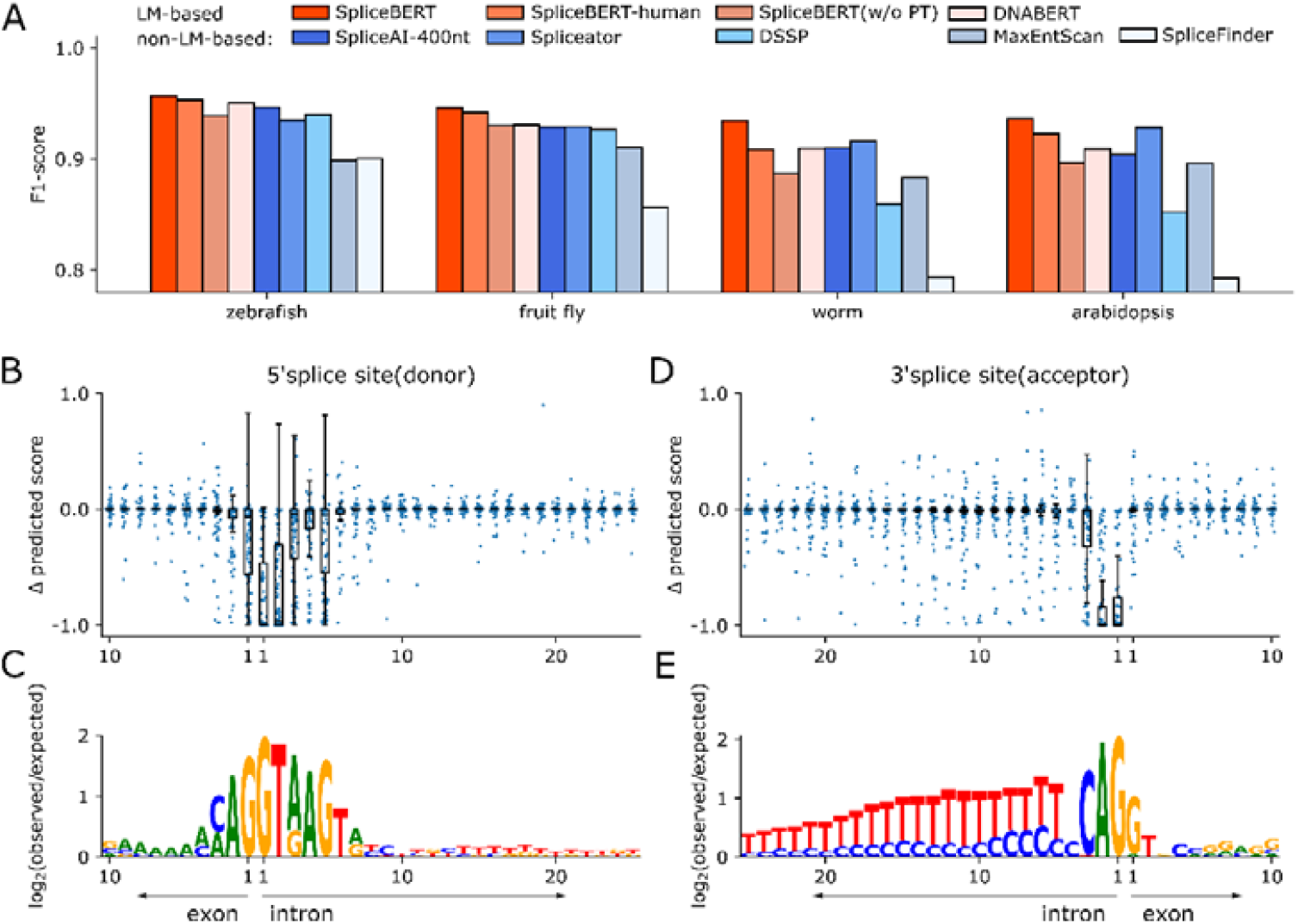
Predicting splice sites in different species using SpliceBERT. (**A**) The F1-score of SpliceBERT and baseline models on 4 species. The visualization of the impact of in silico mutagenesis variants on splice site scores for (**B**) donors and (**D**) acceptors and sequence consensus around splice sites (**C** and **E**).

To confirm the prediction, in silico mutagenesis analysis was performed to compare the putative variant effects with the known pattern of SS. For splice donors, we observed that the variants leading to large change in predicted scores enriched in a 6nt window around donors (1nt in exon and 5nt in intron, Figure 6B). Similarly, for splice acceptors, the variants in the last 3 nucleotides in the 3’end of introns were found to have a substantial impact on predicted scores (Figure 6D). These observations were consistent with the known motif of splice sites^11^, further validating our predictions.

## DISCUSSION

Genetic information, encoded in the genomes, provided a unique challenge for pre-trained language models (pLMs). We are only at the very beginning of unveiling the great learning potential of pLMs in deciphering the complex regulatory logic in genomic sequences. There is still much room for exploration on how to effectively apply pLMs to study genomics. In this study, we focused on RNA splicing and pre-trained the SpliceBERT model on vertebrate pre-mRNA sequences to facilitate the studies of RNA splicing and are particularly interested in how pre-training on vertebrate species will improve the capabilities of pLMs.

By pre-training with masked language modeling, we found that the nucleotide embeddings learned by SpliceBERT not only carried evolutionary conservation information, but can also be utilized to characterize the distinct nature of splice sites. The attention weights in SpliceBERT are able to reflect the associations between splice donors and acceptors from the same introns. Interestingly, we also noticed that splice-disrupting variants tend to induce a larger impact on the embeddings of adjacent nucleotides than benign ones, which was also useful to identify pathogenic variants. With finetuning, SpliceBERT outperformed conventional baseline models as well as human-only pLMs for predicting branchpoints in human and splice sites across species. The improved performance of SpliceBERT may be attributed to the fact that pLMs can generate nucleotide embeddings that are sensitive to sequence context and evolutionary information. These findings demonstrated that self-supervised learning can effectively capture the key determinants of biological function from primary sequences, and this ability can be boosted by integrating a larger amount of pre-training data collected from a diverse range of species.

Currently, SpliceBERT was only pre-trained on vertebrate pre-mRNA sequences, which make up only a small fraction of eukaryotes. More powerful genomic pLMs can be developed by collecting more eukaryotic genomes for pre-training. With the expansion of pre-training dataset, how to improve the quality of dataset will be a critical issue because of the existence of repeats in genomes. As illustrated in our study, MLM on non-repetitive sequences is much more challenging than MLM on repetitive regions. Therefore, reducing the sampling frequency of repetitive regions may improve the training efficiency of genomic pLMs. Besides, SpliceBERT can only process sequences no longer than 1024nt, while prior studies have revealed that leveraging large-scale genomic information can boost the performance of many models in genomics^43–46^, especially for splice sites^10^. Unfortunately, it is not feasible to directly extend SpliceBERT to longer sequences because of the quadratic space complexity of the self-attention model with respect to sequence length. It remains necessary to explore the use of self-attention modules with sub-quadratic complexity^47,48^, or even convolutional networks^49^ for developing more powerful and lightweight genomic language models. Besides, it is also of interest to investigate how to improve the quality of pre-training data, as about half of the genomic sequences are repetitive sequences that of low complexity. More importantly, many biological processes, including RNA splicing, usually occurs in a tissue/cell-specific manner^50^, and sequence-based tissue-specific prediction has proven to be a very difficult task^11,50^. It is worth further study to assess whether pLMs like SpliceBERT can facilitate overcoming these challenges.

## METHODS

### Pre-training SpliceBERT on pre-mRNA sequences

#### Pre-training dataset

We collected pre-mRNA sequences from 72 vertebrates for pre-training. The reference genomes and gene annotations were downloaded from UCSC genome browser^51^, where the genome assembly versions are listed in Table S3. Pre-mRNA sequences were extracted from the reference genomes using “bedtools getfasta”^52^ based on gene annotations and overlapping transcripts were merged to avoid redundancy. In this way, we constructed a pre-training dataset including a total number of about 2 million pre-mRNA sequences that approximately cover 65 billion nucleotides. We reserved 50000 randomly selected sequences for validation and pre-trained SpliceBERT on the remaining sequences.

#### Model architecture

SpliceBERT is based on the BERT^16^ architecture, consisting of 6 Transformer^25^ encoder layers. The hidden layer size and the attention head number are 512 and 16, respectively. Positional information is encoded by the absolution position embedding method and the maximum length of input sequence is set to 1024. SpliceBERT has a total of about 19.4 million trainable parameters.

#### Tokenization

Existing language models for genome sequence mainly adopted the k-mer^22,24,53^ or the Byte Pair Encoding (BPE) tokenization^23^ methods. However, in this study, we simply encoded each nucleotide (A, G, C, T/U) as a token for the ease of obtaining hidden states and attention weights of individual nucleotides. A “[CLS]” (classification) token and a “[SEP]” (separator) token were padded to the beginning and the end of the sequence input to SpliceBERT, respectively, as a routine operation in BERT models^16^.

#### Masked language modeling

We pre-trained SpliceBERT by masked language modeling (MLM), which is a self-supervised learning approach. Specifically, for each input sequence, 15% of the nucleotides were randomly selected. 80% and 10% of the selected nucleotides were replaced with the mask token (“[MASK]”) and random nucleotides, respectively, and the rest of the selected nucleotides remained unchanged. SpliceBERT was trained to predict the correct type of the selected nucleotides during pre-training, by which it will learn the dependencies between the nucleotides and capture the logic of mRNA sequences.

#### Model implementation and pre-training

SpliceBERT was implemented in PyTorch (v1.9) with the Huggingface transformer library (v4.24.0). We adopted the cross entropy loss and the AdamW^54^ optimizer to update the weights in the model. The initial learning rate was set to 0.0001 and will be halved when the loss on the validation data stopped decreasing for 3 epochs. The model was trained in automatic mixed precision mode to reduce memory consumption and speed up model training. We pre-trained the model in two stages. In the first stage, SpliceBERT was pre-trained with sequences of a fixed length of 510nt (512nt with “[CLS]” and “[SEP]”), which were randomly drawn from full-length pre-mRNA sequences. In the second stage, it was pre-trained with sequences of variable length between 64nt and 1024nt (see Supplementary Methods for details). The first stage took 7 days on 8 NVIDIA V100 graphics processing units (GPUs) and the second stage took 3 days on 4 NVIDIA V100 GPUs. The cross-entropy loss on validation data finally converged to 1.02. Additionally, to assess the contribution of multi-species pre-training, we developed another model with identical architecture to SpliceBERT but was pre-trained on only human pre-mRNA sequences, namely SpliceBERT-human.

### The functional genic region, repetitive sequence and evolutionary conservation annotation

For functional genic region annotation, we annotated protein-coding and long non-coding RNA (lncRNA) transcripts against the GENCODE gene annotation (v41lift37, GRCh37/hg19). In protein-coding genes, the genic regions include 5 prime untranslated regions (5’UTR), coding sequences (CDS), 3 prime untranslated regions (3’UTR) and introns. In lncRNA genes, the genic regions include exons and introns. Due to the existence of gene isoforms, the same locus may be annotated into multiple categories. For repetitive sequence (repeat) annotation, we downloaded the RepeatMasker annotation (hg19) in bed format from the UCSC Table browser, and employed “bedtools intersect” to count the proportion of repeats in different functional genic regions. For evolutionary conservation annotation, we downloaded the phastCons^26^ and phyloP^55^ conservation scores derived from multiple sequence alignments of 99 vertebrate genomes to human genome (hg19) from UCSC. The conservation scores can be extracted from bigwig files using the pyBigWig^56^ package.

### Distinguishing conserved/non-conserved sites with nucleotide embedding

Nucleotide embeddings are represented by the hidden states in the last Transformer encoder layer by default. We trained logistic regression (LR) models that took nucleotide embeddings as input to distinguish between nucleotides at conserved (phastCons ≥ 0.8) and non-conserved (phastCons < 0.8) sites. Specifically, the embeddings of nucleotides from 1000 randomly selected 510nt sequences were generated by SpliceBERT, and randomly split into a training dataset (80%) and a test dataset (20%). The nucleotides with unknown (“NaN”) phastCons scores were excluded. Then, an LR model was fitted on the training data and then evaluated on the test dataset. To be noted, no validation dataset was needed here because we simply adopted the default configuration to train LR models. SpliceBERT-human and one-hot encoding were used as baseline methods for comparison. SpliceBERT-human is the model with the same structure as SpliceBERT but was pre-trained on only human pre-mRNA sequences. One-hot encoding is to encode each nucleotide plus its 250nt up- and downstream (501nt in total) with one-hot encodings (A: [1, 0, 0, 0], C: [0, 1, 0, 0], G: [0, 0, 1, 0], T: [0, 0, 0, 1], N: [0, 0, 0, 0]) and flatten into a 2004-dimension vector.

### Visualizing and clustering splice site embeddings

We investigated whether nucleotide embeddings generated by SpliceBERT can be utilized to characterize the nature of splice sites. To this end, splice sites (positive samples) and non-splice sites (negative samples) were collected from the primary transcripts (transcripts with “MANE_Selected” or “Ensembl_canonical” tag) in GENCODE annotation (v41lift37). The positive samples can be directly extracted from gene annotations. As for negative samples, we scanned the transcripts with MaxEntScan^1^ and selected only the non-splice sites with a high MaxEntScan score (>3)^57^ to make it challenging to differentiate between positive and negative samples. The samples that do not match the canonical donor motif GT or acceptor motif AG(U) were discarded to eliminate the difference in nucleotide composition. Finally, we randomly sampled 5000 canonical donor, canonical acceptor, non-donor GT and non-acceptor AG sites, respectively, and generated the embeddings of these samples by SpliceBERT. The embedding of each GT/AT site was concatenated into a 1024-d vector, and then reduced to 128-d by principal component analysis (PCA) for UMAP^27^ visualization. In addition, the Leiden algorithm was performed on the 128-d embeddings to cluster the samples and results were measured by the normalized mutual information (NMI) metric. Here, the programming interfaces of PCA, UMAP and Leiden algorithms are provided in the Scanpy package (v1.9) ^58^. For comparison, we also performed conducted the same analysis on the embeddings generated by SpliceBERT-human, DNABERT and one-hot encoding, respectively. To be noted, the value of k in DNABERT’s tokenization and the size of flanking sequences around splice/non-splice sites impact clustering and visualization, as shown in Supplementary Figure S1 and S2.

### Estimating splice site strength from RNA-seq samples

SpliSER (v0.1.7) ^29^ was utilized to assess the splice site strength (“splice site strength estimation”, SSE) in RNA-seq samples of the K562 cell line. The alignment files (in bam format, GRCh38/hg38) of reversely stranded polyA plus RNA-seq or total RNA-seq were downloaded from the ENCODE project^59,60^ (accession IDs listed in Supplementary Table S4). We removed unmapped or multi-mapped reads (MAPQ < 255) ^13^ from the alignment files by samtools (v1.10)^61^ and then calculated the SSE of splice sites by regtools (v0.5.2)^62^ and SpliSER. Only the splice sites identified in at least 3 samples were reserved and the median SSE value was taken to annotate the splice sites included in the GENCODE annotation (v41, GRCh38/hg38).

### Analyzing attention weights between splice sites

The self-attention module in each Transformer layer of SpliceBERT transforms the input feature map (***h*** ∈ *R*^*L×d*^ of a sequence into a key and a query 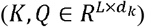 map with two learnable matrices 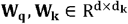 respectively:

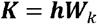

where *d*_*k*_ equals to *d/h*. The dot product of Q and K is the attention matrix:

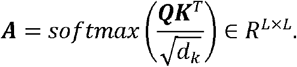

The softmax function normalized the attention matrix by row to make the attention weights in each row sum up to 1. For simplicity, the maximum attention weights across different attention heads were taken. Each element *a*_*ij*_ in A could be regarded as the association of the *j*-th token to the *i*-th token (note that *a*_*ij*_ is not necessarily equal to *a*_*ji*_). In our analysis, we averaged *a*_*ij*_ and *a*_*ji*_ to represent the attention between *i* and *j*. When analyzing the attention weights of donor and acceptor sites, we took the average weights of the two nucleotides in intron and the one nucleotide in exon.

### Measuring the impact of genetic variants using KL-divergence

In masked language modeling, SpliceBERT gives a nucleotide type probability distribution for each token based on sequence context. We leveraged this feature to estimate the impact of variants on RNA splicing in an unsupervised manner. Intuitively, the nucleotide type distribution of the *i*th nucleotide in a sequence without variants can be represented as P_*i*_ *=* (*p*_*A*_, *p*_*C*_, *p*_*G*_, *p*_*T*_), where *p*_*A*_, *p*_*C*_, *p*_*G*_ sums *p*_*T*_ up to 1. When a variant occurs in the sequence, it will introduce perturbations to the model’s output, resulting in the probability distribution of the nucleotide *i* change from 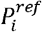 to 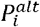. Then, the change of distribution can be measured by Kullback-Leibler (KL) divergence:

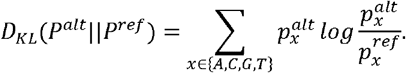

The probability values were clipped to be between [10^−6^, 1] to avoid division by zero error. We recruited two datasets of variants associated with RNA splicing (MFASS and Vex-seq) to illustrate the effectiveness of our approach. The MFASS dataset includes 27733 single nucleotide variants (SNVs) within or around exons. The SNVs that largely decrease exon splicing efficiency (ΔΨ < -0.5, Ψ: PSI/Percent Spliced In, n = 1050) were considered as splice-disrupting variants^34^. The Vex-seq dataset includes 1971 SNVs and 84 indels and the ΔΨ included by the variants. We recruited only the SNVs and defined the variants with ΔΨ < −0.24 (top 5%, n = 98) as SDVs.

### In silico mutagenesis analysis

In silico mutagenesis (ISM) ^63^ was employed to investigate the effects of genetic variants on their neighboring nucleotides or branchpoint/splice site predictions. Specifically, we enumerated all possible single nucleotide variants (SNVs) in whole or a custom subregion of each input sequence. Only a single SNV is introduced to a sequence each time. The sequences without and with variants are referred to as wild-type (WT) and mutant (MT) sequences, respectively. The difference between the predictions given by the model for MT and WT sequences can be used to estimate the effects of variants.

### Finetuning SpliceBERT for predicting branchpoint

SpliceBERT was finetuned on a human branchpoint (BP) dataset^40^ (the Mercer’s dataset) for predicting branchpoint (BP) sites. The Mercer’s dataset includes branchpoints of high confidence (HC-BP) and low confidence (LC-BP) identified in the human transcripts. Following previous studies^40,64^, we focused on the 18-44nt region upstream of splice acceptors (where BPs typically located) and recruited the HC-BP (n = 55739) and non-BP (n = 921518) sites within the focused regions as positive and negative samples, respectively. The input to SpliceBERT is a 510nt sequence that covers both the up- and downstream of the focused regions, and the hidden states in the last Transformer layer of SpliceBERT were fed to a single-layer fully-connected neural network to predict BPs (implemented by the “BertForTokenClassification” class in the Huggingface transformers package). To improve prediction efficiency, multiple sites in a single sequence can be predicted by SpliceBERT simultaneously. The nested cross-validation (CV) strategy was employed to finetune and evaluate SpliceBERT. The samples were split into 10 folds by chromosomes such that the samples from the same chromosome were always kept in the same CV fold. In each training epoch, each of the 10 CV folds was reserved in turn as the test dataset, and the other folds were used to finetune and validate SpliceBERT (8 folds for training and 1 fold for validating). The optimal number of training epochs was determined according to the average performance on the 10 folds for validation and the training process will be terminated when the CV performance stopped to improve for 10 epochs. The final performance of the model is measured by the average precision (AP) score on the 10 test CV folds. We compared SpliceBERT to Branchpointer^39,40^, LaBranchoR, DNABERT and SpliceBERT-human. The predictions of Branchpointer were generated using its scripts and the results of other models were obtained by training and testing them using the same nested CV scheme to SpliceBERT.

### Finetuning SpliceBERT for predicting splice sites

SpliceBERT was finetuned on the Spliceator dataset^42^ to predict splice sites across species. The Spliceator dataset curated confirmed error-free splice sites from over 100 eukaryotic species and provided 5 independent test datasets from vertebrates (*Homo sapiens* – human, *Danio rerio* – zebrafish), invertebrates (*Drosophila melanogaster* – fruit fly, *Caenorhabditis elegans* – worm) and plants (*Arabidopsis thaliana* – arabidopsis). Because human splice sites are also included in training data, we recruited only the 4 non-human test datasets to evaluate the model’s ability for cross-species splice site prediction. Each sample in the dataset is a 600nt/400nt sequence centered on a splice/non-splice site. To ensure the consistency of all samples, the sequences of 600nt were truncated to 400nt. We finetuned SpliceBERT as a sequence classification task, feeding the hidden state of the “[CLS]” token in the last layer to a two-layer fully-connected neural network to make predictions. To make our results comparable to those reported in the Spliceator paper, we followed the 10-fold cross-validation scheme described in the study to finetune SpliceBERT and test it on the same independent datasets. Baseline models for comparison include SpliceBERT-human, DNABERT, SpliceAI-400nt, Spliceator, DSSP, MaxEntScan and SpliceFinder. The results (measure by the F1 score) of Spliceator, DSSP, MaxEntScan and SpliceFinder were directly taken from the Spliceator paper, and the results of the other models were obtained by training and testing them following the same procedure we used to finetune SpliceBERT. To be noted, the state-of-the-art SpliceAI^10^ model (SpliceAI-10k) takes ultra-long sequences of at least 10001nt, which exceeds the maximum length that our model can process (L=1024nt). Thus, we only compared SpliceAI-400nt with other models.

### Visualization of consensus sequences

We visualized the consensus sequences around branchpoints and splice sites using the Logomaker package ^65^. The logarithm of the ratio between observed and expected nucleotide frequency was taken for visualization: 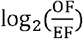. The observed frequencies (OF) at each position were obtained by taking the average occurrence of nucleotide types across different sequences. The expected frequency (EF) of each nucleotide type (background frequency) was obtained by taking the average occurrence frequency of entire sequences.

## AVAILABILITY

The source code and model weights of SpliceBERT are available at https://github.com/biomed-AI/SpliceBERT. The reference genomes and gene annotations of vertebrates are at https://hgdownload.soe.ucsc.edu/downloads.html. The GENCODE annotation is at https://www.gencodegenes.org/human/release_41lift37.html. The annotation of repeats was obtained through the UCSC Table Browser https://genome.ucsc.edu/cgi-bin/hgTables. PhastCons and phyloP conservation scores are at http://hgdownload.cse.ucsc.edu/goldenpath/hg19/phastCons100way/ and http://hgdownload.cse.ucsc.edu/goldenpath/hg19/phyloP100way/. The RNA-seq samples of K562 are at https://www.encodeproject.org/, with accession IDs listed in Supplementary Table S3.

## REFERENCES

1. Yeo, G. & Burge, C. B. Maximum entropy modeling of short sequence motifs with applications to RNA splicing signals. J. Comput. Biol. 11, 377–394 (2004).

2. Barash, Y. et al. Deciphering the splicing code. Nature 465, 53 (2010).

3. Xiong, H. Y. et al. The human splicing code reveals new insights into the genetic determinants of disease. Science 347, 1254806–1254806 (2015).

4. Mount, S. M. et al. Assessing predictions of the impact of variants on splicing in CAGI5. Human Mutation 40, 1215–1224 (2019).

5. Cartegni, L., Wang, J., Zhu, Z., Zhang, M. Q. & Krainer, A. R. ESEfinder: A web resource to identify exonic splicing enhancers. Nucleic Acids Res 31, 3568–3571 (2003).

6. Zhang, Q. et al. BPP: a sequence-based algorithm for branch point prediction. Bioinformatics 33, 3166–3172 (2017).

7. Desmet, F.-O. et al. Human Splicing Finder: an online bioinformatics tool to predict splicing signals. Nucleic Acids Research 37, e67–e67 (2009).

8. Rosenberg, A. B., Patwardhan, R. P., Shendure, J. & Seelig, G. Learning the Sequence Determinants of Alternative Splicing from Millions of Random Sequences. Cell 163, 698–711 (2015).

9. Cheng, J. et al. MMSplice: modular modeling improves the predictions of genetic variant effects on splicing. Genome Biology 20, 48 (2019).

10. Jaganathan, K. et al. Predicting Splicing from Primary Sequence with Deep Learning. Cell 176, 535–548.e24 (2019).

11. Zeng, T. & Li, Y. I. Predicting RNA splicing from DNA sequence using Pangolin. Genome Biology 23, 103 (2022).

12. Trapnell, C. et al. Transcript assembly and quantification by RNA-Seq reveals unannotated transcripts and isoform switching during cell differentiation. Nat Biotechnol 28, 511–515 (2010).

13. Dobin, A. et al. STAR: ultrafast universal RNA-seq aligner. Bioinformatics 29, 15–21 (2012).

14. Gao, K., Masuda, A., Matsuura, T. & Ohno, K. Human branch point consensus sequence is yUnAy. Nucleic Acids Research 36, 2257–2267 (2008).

15. Mikolov, T., Sutskever, I., Chen, K., Corrado, G. S. & Dean, J. Distributed Representations of Words and Phrases and their Compositionality. in Advances in Neural Information Processing Systems 26 (eds. Burges, C. J. C., Bottou, L., Welling, M., Ghahramani, Z. & Weinberger, K. Q.) 3111–3119 (Curran Associates, Inc., 2013).

16. Devlin, J., Chang, M.-W., Lee, K. & Toutanova, K. BERT: Pre-training of Deep Bidirectional Transformers for Language Understanding. in Proceedings of the 2019 Conference of the North American Chapter of the Association for Computational Linguistics: Human Language Technologies, Volume 1 (Long and Short Papers) 4171–4186 (Association for Computational Linguistics, 2019). doi:10.18653/v1/N19-1423.

17. Radford, A., Narasimhan, K., Salimans, T. & Sutskever, I. Improving Language Understanding by Generative Pre-Training.

18. Rives, A. et al. Biological structure and function emerge from scaling unsupervised learning to 250 million protein sequences. Proceedings of the National Academy of Sciences 118, e2016239118 (2021).

19. Elnaggar, A. et al. ProtTrans: Towards Cracking the Language of Lifes Code Through Self-Supervised Deep Learning and High Performance Computing. IEEE Transactions on Pattern Analysis and Machine Intelligence 1–1 (2021) doi:10.1109/TPAMI.2021.3095381.

20. Chen, J. et al. Interpretable RNA Foundation Model from Unannotated Data for Highly Accurate RNA Structure and Function Predictions. Preprint at https://doi.org/10.48550/arXiv.2204.00300 (2022).

21. Zvyagin, M. et al. GenSLMs: Genome-scale language models reveal SARS-CoV-2 evolutionary dynamics. http://biorxiv.org/lookup/doi/10.1101/2022.10.10.511571 (2022) doi:10.1101/2022.10.10.511571.

22. Yang, M. et al. Integrating convolution and self-attention improves language model of human genome for interpreting non-coding regions at base-resolution. Nucleic Acids Research gkac326 (2022) doi:10.1093/nar/gkac326.

23. Cahyawijaya, S. et al. SNP2Vec: Scalable Self-Supervised Pre-Training for Genome-Wide Association Study. in Proceedings of the 21st Workshop on Biomedical Language Processing 140–154 (Association for Computational Linguistics, 2022). doi:10.18653/v1/2022.bionlp-1.14.

24. Ji, Y., Zhou, Z., Liu, H. & Davuluri, R. V. DNABERT: pre-trained Bidirectional Encoder Representations from Transformers model for DNA-language in genome. Bioinformatics 37, 2112–2120 (2021).

25. Vaswani, A. et al./person-group>. Attention is All you Need. in Advances in Neural Information Processing Systems 30 (eds. Guyon, I. et al.) 5998–6008 (Curran Associates, Inc., 2017).

26. Siepel, A. et al. Evolutionarily conserved elements in vertebrate, insect, worm, and yeast genomes. Genome Res 15, 1034–1050 (2005).

27. McInnes, L., Healy, J. & Melville, J. UMAP: Uniform Manifold Approximation and Projection for Dimension Reduction. arXiv:1802.03426 [cs, stat] (2020).

28. Traag, V. A., Waltman, L. & van Eck, N. J. From Louvain to Leiden: guaranteeing well-connected communities. Sci Rep 9, 5233 (2019).

29. Dent, C. I. et al. Quantifying splice-site usage: a simple yet powerful approach to analyze splicing. NAR Genomics and Bioinformatics 3, lqab041 (2021).

30. Rogers, A., Kovaleva, O. & Rumshisky, A. A Primer in BERTology: What We Know About How BERT Works. Transactions of the Association for Computational Linguistics 8, 842–866 (2020).

31. Tilgner, H. et al. Deep sequencing of subcellular RNA fractions shows splicing to be predominantly co-transcriptional in the human genome but inefficient for lncRNAs. Genome Res. 22, 1616–1625 (2012).

32. Li, Y. I. et al. Annotation-free quantification of RNA splicing using LeafCutter. Nature Genetics 50, 151 (2018).

33. Kalmykova, S. et al. Conserved long-range base pairings are associated with pre-mRNA processing of human genes. Nat Commun 12, 2300 (2021).

34. Cheung, R. et al. A Multiplexed Assay for Exon Recognition Reveals that an Unappreciated Fraction of Rare Genetic Variants Cause Large-Effect Splicing Disruptions. Molecular Cell 73, 183–194.e8 (2019).

35. Adamson, S. I., Zhan, L. & Graveley, B. R. Vex-seq: high-throughput identification of the impact of genetic variation on pre-mRNA splicing efficiency. Genome Biology 19, 71 (2018).

36. Meier, J. et al. Language models enable zero-shot prediction of the effects of mutations on protein function. in Advances in Neural Information Processing Systems vol. 34 29287–29303 (Curran Associates, Inc., 2021).

37. Benegas, G., Batra, S. S. & Song, Y. S. DNA language models are powerful zero-shot predictors of non-coding variant effects. 2022.08.22.504706 Preprint at https://doi.org/10.1101/2022.08.22.504706 (2022).

38. Mercer, T. R. et al. Genome-wide discovery of human splicing branchpoints. Genome Res 25, 290–303 (2015).

39. Paggi, J. M. & Bejerano, G. A sequence-based, deep learning model accurately predicts RNA splicing branchpoints. RNA 24, 1647–1658 (2018).

40. Signal, B., Gloss, B. S., Dinger, M. E. & Mercer, T. R. Machine learning annotation of human branchpoints. Bioinformatics 34, 920–927 (2018).

41. Zhang, P. et al. Genome-wide detection of human variants that disrupt intronic branchpoints. Proceedings of the National Academy of Sciences 119, e2211194119 (2022).

42. Scalzitti, N. et al. Spliceator: multi-species splice site prediction using convolutional neural networks. BMC Bioinformatics 22, 561 (2021).

43. Schwessinger, R. et al. DeepC: predicting 3D genome folding using megabase-scale transfer learning. Nat Methods 17, 1118–1124 (2020).

44. Chen, K., Zhao, H. & Yang, Y. Capturing large genomic contexts for accurately predicting enhancer-promoter interactions. Briefings in Bioinformatics 23, bbab577 (2022).

45. Avsec, Ž. et al. Effective gene expression prediction from sequence by integrating long-range interactions. Nat Methods 18, 1–8 (2021).

46. Lee, D., Yang, J. & Kim, S. Learning the histone codes with large genomic windows and three-dimensional chromatin interactions using transformer. Nat Commun 13, 6678 (2022).

47. Choromanski, K. et al. Rethinking Attention with Performers. arXiv:2009.14794 [cs, stat] (2020).

48. Hua, W., Dai, Z., Liu, H. & Le, Q. Transformer Quality in Linear Time. in Proceedings of the 39th International Conference on Machine Learning 9099–9117 (PMLR, 2022).

49. Yang, K. K., Fusi, N. & Lu, A. X. Convolutions are competitive with transformers for protein sequence pretraining. 2022.05.19.492714 Preprint at https://doi.org/10.1101/2022.05.19.492714 (2022).

50. Cheng, J., Çelik, M. H., Kundaje, A. & Gagneur, J. MTSplice predicts effects of genetic variants on tissue-specific splicing. Genome Biology 22, 94 (2021).

51. Haeussler, M. et al. The UCSC Genome Browser database: 2019 update. Nucleic Acids Res. 47, D853–D858 (2019).

52. Quinlan, A. R. & Hall, I. M. BEDTools: a flexible suite of utilities for comparing genomic features. Bioinformatics 26, 841–842 (2010).

53. Ng, P. dna2vec: Consistent vector representations of variable-length k-mers. arXiv:1701.06279 [cs, q-bio, stat] (2017).

54. Loshchilov, I. & Hutter, F. Decoupled Weight Decay Regularization. Preprint at https://doi.org/10.48550/arXiv.1711.05101 (2019).

55. Pollard, K. S., Hubisz, M. J., Rosenbloom, K. R. & Siepel, A. Detection of nonneutral substitution rates on mammalian phylogenies. Genome Res 20, 110–121 (2010).

56. Ramírez, F. et al. deepTools2: a next generation web server for deep-sequencing data analysis. Nucleic Acids Res 44, W160–W165 (2016).

57. Bretschneider, H., Gandhi, S., Deshwar, A. G., Zuberi, K. & Frey, B. J. COSSMO: predicting competitive alternative splice site selection using deep learning. Bioinformatics 34, i429–i437 (2018).

58. Wolf, F. A., Angerer, P. & Theis, F. J. SCANPY: large-scale single-cell gene expression data analysis. Genome Biology 19, 15 (2018).

59. ENCODE Project Consortium. An integrated encyclopedia of DNA elements in the human genome. Nature 489, 57–74 (2012).

60. Davis, C. A. et al. The Encyclopedia of DNA elements (ENCODE): data portal update. Nucleic Acids Res. 46, D794–D801 (2018).

61. Li, H. et al. The Sequence Alignment/Map format and SAMtools. Bioinformatics 25, 2078–2079 (2009).

62. Cotto, K. C. et al. Integrated analysis of genomic and transcriptomic data for the discovery of splice-associated variants in cancer. Nat Commun 14, 1589 (2023).

63. Zhou, J. & Troyanskaya, O. G. Predicting effects of noncoding variants with deep learning–based sequence model. Nat Methods 12, 931–934 (2015).

64. Avsec, Ž., Barekatain, M., Cheng, J. & Gagneur, J. Modeling positional effects of regulatory sequences with spline transformations increases prediction accuracy of deep neural networks. Bioinformatics 34, 1261–1269 (2018).

65. Tareen, A. & Kinney, J. B. Logomaker: beautiful sequence logos in Python. Bioinformatics 36, 2272–2274 (2020).

