## Supplementary Figure for "Self-supervised learning on millions of pre-mRNA sequences improves sequence-based RNA splicing prediction"

Supplementary Materials

Ken Chen[1], Yue Zhou[2], Maolin Ding[1], Yu Wang[2], Zhixiang Ren[2,*], and Yuedong Yang[1,3,*]

[1]School of Computer Science and Engineering, Sun Yat-sen University, Guangzhou, Guangdong, China

[2]Pengcheng Laboratory, Shenzhen, Guangdong, China

[3]Key Laboratory of Machine Intelligence and Advanced Computing (Sun Yat-sen University), Ministry of Education, China

### 1 Pre-training SpliceBERT

We downloaded the latest reference genomes and gene annotations of vertebrates from the USCS Genome Browser[1] (Haeussler et al., 2019) in Jul. 2022, where the versions are available in Table S1. The genome files in uncompressed fasta format exceeds 150 GB and cannot be loaded into memory entirely. Therefore, we converted each chromosome to a numpy array (data type: `numpy.int8`) with one-hot encoding (N: 0, A: 1, C: 2, G: 3, T: 4) and saved them in `hdf5` format. Then, the transcript sequences can be randomly accessed from disk with genomic coordinates during model training. For pre-training, we sampled transcripts in proportion to their sequence length. Since the length of transcripts varies greatly and more than 80 % are longer than 1024nt, sequences fragments no longer than 1024nt were randomly drawn from full-length transcripts. When the selected fragments exceeded the boundary of the transcript, the flanking genomic sequences of the transcript will be padded. The sampling process was performed on the fly during pre-training, which means that samples used for pre-training were different in each epoch.

SpliceBERT was pre-trained for two stages. The first stage was pre-trained on fixed-length sequences of 510nt. Since SpliceBERT uses learnable position embeddings to preserve positional information, it can not be directly applied to sequences longer than 510nt. To make the model converge faster, we adopted a hierarchical decomposition strategy proposed by (Su, 2020) to extend the positional embeddings to a maximum of 1024nt. Briefly, we use $\boldsymbol{p}_1, \boldsymbol{p}_2, \cdots, \boldsymbol{p}_m, \boldsymbol{p}_{m+1}, \cdots, \boldsymbol{p}_n$ to represent the position embeddings in the model (e.g., $m = 510, n = 1024$ in our study), where $\boldsymbol{p}_1, \cdots, \boldsymbol{p}_m$ have been trained in the first stage. The hierarchical decomposition strategy decomposes the embeddings into:

$$\boldsymbol{p}_{(i-1)\times m+j} = \alpha \boldsymbol{u}_i + (1-\alpha)\boldsymbol{u}_j, \qquad i, j \in \{1, 2, \cdots, m\} \tag{1}$$

, where $\boldsymbol{u}_i = \dfrac{\boldsymbol{p}_i - \alpha \boldsymbol{p}_1}{1-\alpha}, \alpha \neq 0.5$ ($\alpha = 0.4$ by default). This strategy can speed up the pre-training process in the second stage.

### References

Haeussler, M., Zweig, A. S., Tyner, C., Speir, M. L., Rosenbloom, K. R., Raney, B. J., … Kent, W. J. (2019). The UCSC Genome Browser database: 2019 update. *Nucleic Acids Res.*, *47*(D1), D853–D858. doi: 10.1093/nar/gky1095

Su, J. (2020, Dec). *Hierarchical decomposition of positional encoding enables BERT to handle longer sequences (in Chinese).* Retrieved from `https://spaces.ac.cn/archives/7947`

[1] `https://hgdownload.cse.ucsc.edu/goldenpath/`

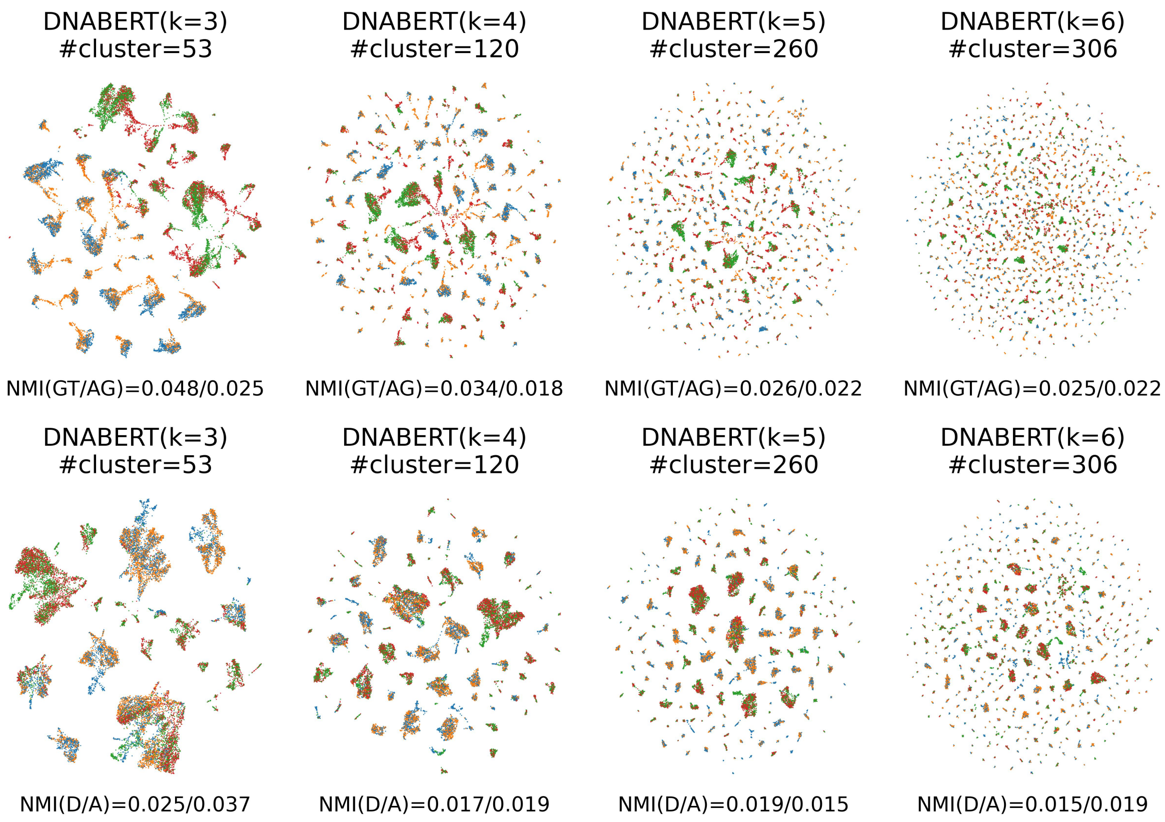

Figure S1: Comparison of UMAP visualization of embeddings of (up) splice/non-spilce sites and (down) strong/weak splice sites obtained by DNABERT with different token lengths (k=3, 4, 5, 6). The Leiden algorithm was employed to cluster the embeddings.

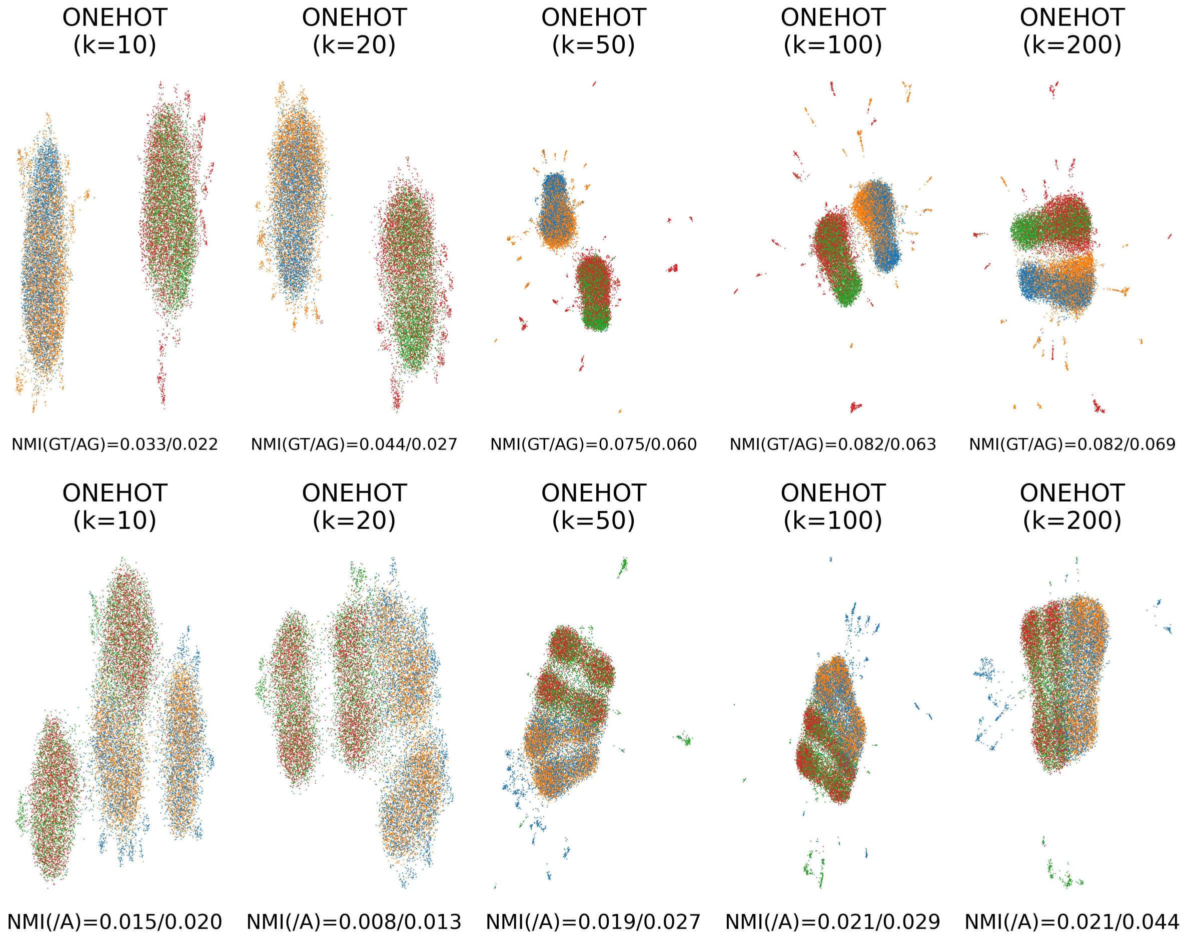

Figure S2: UMAP of (**up**) splice/non-splice site and (**down**) strong/weak splice site embeddings acquired by one-hot encoded sequences of different length. The Leiden algorithm was employed to cluster the embeddings.

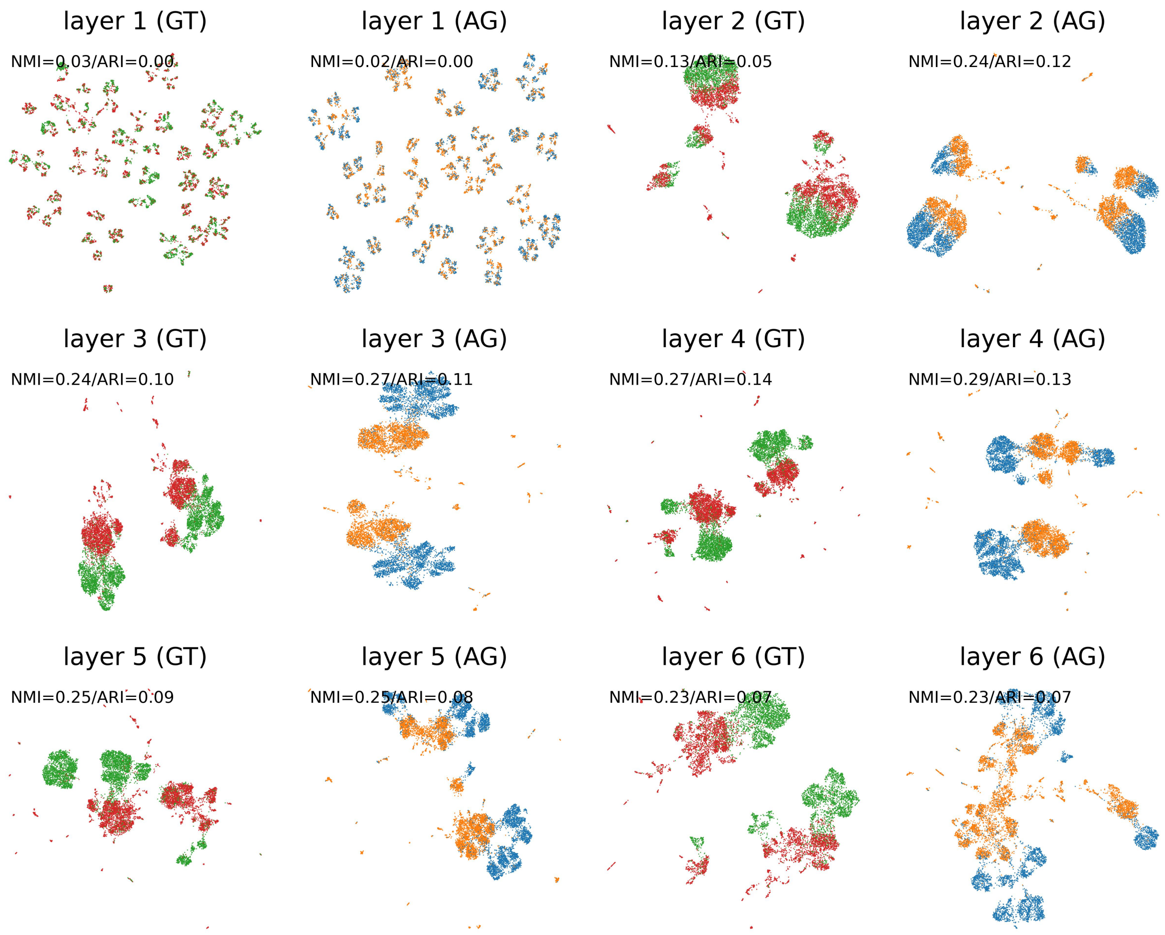

Figure S3: Comparison of nucleotide embeddings from different Transformer encoder layers

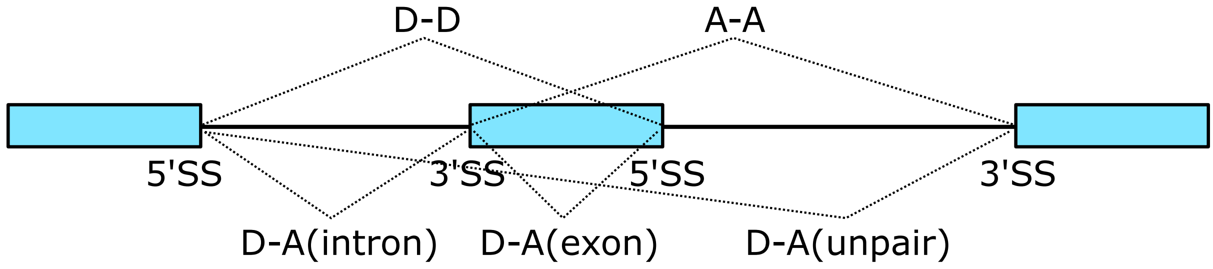

Figure S4: Different combination of donor/acceptor sites

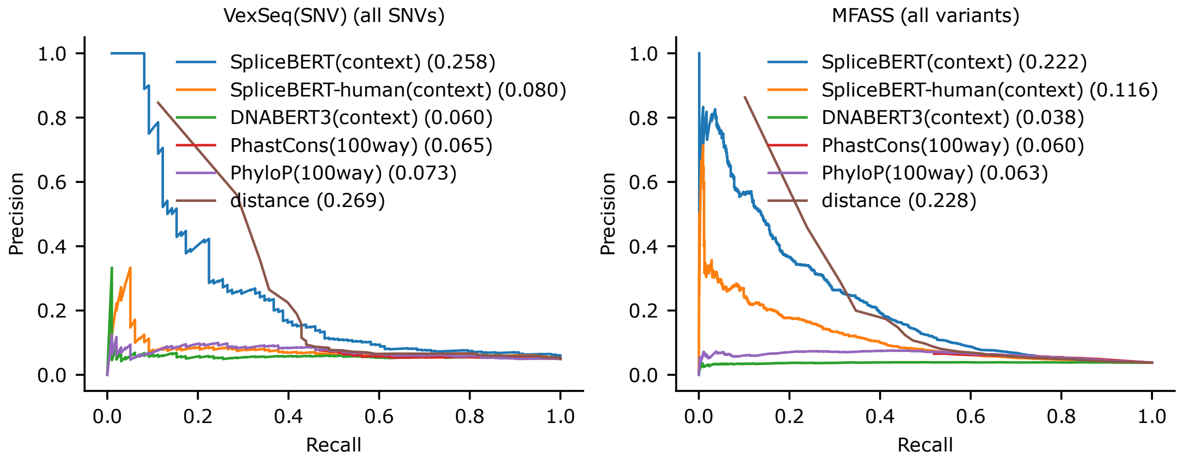

Figure S5: Precision-recall curves of SpliceBERT and baseline methods for unsupervised variants effect prediction. When all variants are considered, the distance (distance from splice sites to variants) dominates the performance because variants reside in splice sites are much more likely to affect splicing.
